# *Prunella vulgaris* extract and suramin block SARS-coronavirus *2* virus Spike protein D614 and G614 variants mediated receptor association and virus entry in cell culture system

**DOI:** 10.1101/2020.08.28.270306

**Authors:** Zhujun Ao, Mable Chan, Maggie Jing Ouyang, Olukitibi Titus Abiola, Mona Mahmoudi, Darwyn Kobasa, Xiaojian Yao

**Author notes:** To whom correspondence should be addressed to: X-j. Yao, Dept. of Medical Microbiology, Max Rady College of Medicine, Faculty of Health Sciences, University of Manitoba, 508-745 Bannatyne Ave, Winnipeg MB R3E 0J9.

## Abstract

Until now, no approved effective vaccine and antiviral therapeutic are available for treatment or prevention of SARS-coronavirus 2 (SCoV-2) virus infection. In this study, we established a SCoV-2 Spike glycoprotein (SP), including a SP mutant D614G, pseudotyped HIV-1-based vector system and tested their ability to infect ACE2-expressing cells. This study revealed that a C-terminal 17 amino acid deletion in SCoV-2 SP significantly increases the incorporation of SP into the pseudotyped viruses and enhanced its infectivity, which may be helpful in the design of SCoV2-SP-based vaccine strategies. Moreover, based on this system, we have demonstrated that an aqueous extract from the Chinese herb *Prunella vulgaris* (CHPV) and a compound, suramin, displayed potent inhibitory effects on both wild type and mutant (G614) SCoV-2 SP pseudotyped virus (SCoV-2-SP-PVs)-mediated infection. The 50% inhibitory concentration (IC50) for CHPV and suramin on SCoV-2-SP-PVs are 30, and 40 μg/ml, respectively. To define the mechanisms of their actions, we demonstrated that both CHPV and suramin are able to directly interrupt SCoV-2–SP binding to its receptor ACE2 and block the viral entry step. Importantly, our results also showed that CHPV or suramin can efficiently reduce levels of cytopathic effect caused by SARS-CoV-2 virus (hCoV-19/Canada/ON-VIDO-01/2020) infection in Vero cells. Furthermore, our results demonstrated that the combination of CHPV/suramin with an anti-SARS-CoV-2 neutralizing antibody mediated more potent blocking effect against SCoV2-SP-PVs. Overall, this study provides evidence that CHPV and suramin has anti-SARS-CoV-2 activity and may be developed as a novel antiviral approach against SARS-CoV-2 infection.

## Introduction

The recent and ongoing outbreak of Coronavirus disease 2019 (COVID-19) has called for serious and urgent global attention [1,2]. The COVID-19 disease is caused by a newly emerged virus strain of Severe Acute Respiratory Syndrome (SARS) known as SARS-CoV-2 [3]. Although the case fatality ratio (CFR) of COVID-19 can only be detected at the end of the outbreak, an estimated global CFR was calculated to be 5.5-5.7% in March 2020, which is shockingly more than seasonal influenza outbreak [4]. While in August 2020, the infection fatality ratio was estimated by WHO to be 0.5-1% [5]. Since the identification of the SARS-CoV-2 sequences [6], extensive efforts worldwide have been focused on the development of vaccine and antiviral drugs against SARS-CoV-2 with the hope to rapidly and efficiently control this new human coronavirus (CoV) infection.

SARS-CoV-2 belongs to a betacoronavirus subfamily that includes enveloped, large and positive-stranded RNA viruses responsible for causing severe respiratory system, gastrointestinal and neurological symptoms [7-10]. The human CoV was first identified in 1960 and constituted about 30% of the causes of the common cold. Among the identified human CoVs are NL63, 229E, OC43, HKU1, SARS-CoV, the Middle East respiratory syndrome (MERS)-CoV, and SARS-CoV-2 [11,12]. A recent study has revealed that SARS-CoV-2 was closely related (88% identity) to two SARS-like CoVs that were isolated from bats in 2018 in China, but it was less related to SARS-CoV (79%) and MERS-CoV (about 50%) [13]. The key determinant for the infectivity of SARS-CoV-2 depends on the host specificity with the viral surface-located trimeric spike (S) glycoprotein (SP), which is commonly cleaved by host proteases into an N-terminal S1 subunit and a membrane-embedded C-terminal S2 region [14]. Recent studies revealed that a SP mutation, Aspartic acid (D) changed to Glycine (G) at amino acid position 614, in the S1 domain has been found in high frequency (65% to 70%) in April to May of 2020, that was associated with an increased viral load and significantly higher transmission rate in infected individuals, but no significant change with disease severity [15]. Following studies also suggested that G614 SP mutant pseudotyped retroviruses infected ACE2-expressing cells markedly more efficiently than those with D614 SP [16].

Up till now, several compounds have been tested in numerous clinical trials, including remdesivir, lopinavir, umifenovir, and hydroxychloroquine [17-21]. Moreover, some *in vitro* research suggested that other drugs such as fusion peptide (EK1), anti-inflammatory drugs (such as hormones and other molecules) could be potentially used in the treatment of SARS-CoV-2 disease (reviewed in [22,23]). However, their safety and efficacy have not been confirmed by clinical trials. Currently, specific antiviral treatment drugs are still not available for SARS-CoV-2 infections [23].

Traditional Chinese medicine holds a unique position among all conventional medicines because of its usage over more than hundreds of years of history. Many aqueous extracts of traditional Chinese medicinal herbs have been proven to have antiviral activities [24], and most of these are generally of low toxicity, cheap and readily accessible. As an easily accessible and low-cost natural source, they are especially valuable as potential new sources for rapid responses against the ongoing COVID-19 pandemic. *Prunella vulgaris*, widely distributed in China, Europe, northwestern Africa and North America, is known as a self-heal herb and studies have previously found that a water-soluble substance from Chinese Herb *Prunella vulgaris* (CHPV) exhibit significant antiviral activity against HIV, HSV and Ebola virus [25-28]. However, whether CHPV can block SARS-CoV-2 virus infection is unknown. Another compound, Suramin, has also been previously shown to be a potent inhibitor against HIV [29], while the subsequent studies revealed that its inhibitory effects on HIV replication did not correlate with clinical or immunologic improvement [30,31]. A previous study observed that suramin not only substantially reduced viral loads of *chikungunya virus* (CHIKV) in infected mice, but it also ameliorated virus-induced foot lesions in the mice [32]. Recently, Salgado-Benvindo C., *et al*., reported that Suramin is able to inhibit SARS-CoV-2 infection in cell culture by interfering with early steps of the replication cycle [33].

In this study, we have established a highly sensitive SARS-CoV-2 SP-pseudotyped virus (SCoV-2 SP-PVs) system and investigated the impact of the cytoplasmic tail and a G614 mutant of SP on virus entry ability. We also examined two compounds, CHPV and suramin, for their blocking activities in the SCoV-2 SP-PVs system and SARS-CoV-2 infection, and the antiviral mechanism of their actions. Furthermore, we investigated the synergistic effect of combining anti-SARS-CoV-2 neutralizing antibody (nAb) with CHPV or suramin to enhance their anti-SARS-CoV-2 activity. Overall, this study provides evidence for the first time that CHPV, an aqueous extract from *Prunella vulgaris*, has potent anti-SARS-CoV-2 activity.

## Materials and methods

### Plasmid constructs

The SARS-CoV-2 expressing plasmids (pCAGGS-nCoVSP, pCAGGS-nCoVSPΔC and pCAGGS-nCoVSPΔC_G614_) containing SARS-CoV-2 SP transgene (GenBank accession No. MN908947) or corresponding mutated genes for SPΔC and ΔC_G614_. The SPΔC and ΔC_G614,_ were generated by mutagenic PCR technique. Primers are following: SPΔC-3’primer, 5_GCAGGTACCTAGAATTTGCAGCAGGATCCAC; D614G-5’, 5_GCTGTTCTTTATCAGGGTGTTAACTGCACAG; D614G-3’, 5_CTGTGCAGTTAACACCCTGATAAAGAACAGC. Mutated genes were cloned into the pCAGGS plasmid and each mutation was confirmed by sequencing. The HIV vector encoding for Gaussia luciferase gene HIV-1 RT/IN/Env tri-defective proviral plasmid (ΔRI/E/Gluc) and the helper packaging plasmid pCMVΔ8.2 encoding for the HIV Gag-Pol plasmids are described previously [26,34].

### Cell culture, antibodies and chemicals

The human embryonic kidney cells (HEK293T) and kidney epithelial cells (VeroE6 and Vero cells (ATCC, CCL-81)) from African green monkey were cultured in Dulbecco’s modified Eagle’s medium (HEK293T, VeroE6) or Minimum Essential Medium (MEM; Vero). HEK293T expressing ACE2 (293T_ACE2_) was obtained from GeneCopoeia Inc, Rockville, MD. All cell lines were supplemented with 10% fetal bovine serum (FBS), 1X L-Glutamine and 1% penicillin and streptomycin. The rabbit polyclonal antibody against SARS-CoV-2 SP (Cat# 40150-R007) and ACE2 protein (Cat# 40592-T62) were obtained from Sino Biological and anti-HIVp24 monoclonal antibody was described previously [35,36]. The HIV-1 p24 ELISA Kit was obtained from the AIDS Vaccine Program of the Frederick Cancer Research and Development Center. SARS-CoV-2 SP-ACE2 binding ELISA kit (Cat# COV-SACE2-1) was purchased from RayBio. Anti-SARS-CoV-2 neutralizing Antibody (nAb) Human IgG1(SAD-535) was purchased from ACRO Biosystems..

### Preparation and purification of herb extracts of P. vulgaris L (CHPV)

The dried fruitspikes of *P. vulgaris L*. (Labiatae) (Fig. 3A) were first soaked overnight in deionized water at room temperature and then boiled for one hour. Then the cooled supernatant was centrifuged (3000 g, 30 min), filtered through a 0.45 μm cellulose acetate membrane and finally lyophilized, as described previously [25]. The resulting dark brown residue was dissolved in deionized water and stored at −20°C. A single symmetrical peak corresponding to a molecular weight of polysaccharides (approximately 10 kDa) in the aqueous extract from PV was detected by HPLC analysis, as described previously [25]. Suramin (Cat# sc-200833) was purchased from Santa Cruz BioTech and was dissolved in sterile H2O and stored at −20°.

### Virus production, infection and inhibition experiments

SARS-CoV-2 SP or SPΔC pseudotyped viruses (SCoV-2-SP-PVs, SCoV-2-SPΔC-PVs, SCoV-2-SPΔC_G614_-PVs) were produced by transfecting HEK293T cells with pCAGGS-SARS-CoV-2-SP, pCAGGS-SARS-CoV-2-SPΔC, or pCAGGS-SARS-CoV-2-SPΔC_G614_, pCMVΔ8.2 and a Gluc expressing HIV vector ΔRI/E/Gluc. After 48 hrs of transfection, cell culture supernatants were collected and pseudotyped VLPs were purified from the supernatant by ultracentrifugation (32,000 rpm) for 2 hrs. The pelleted VPs were resuspended into RPMI medium and virus titers were quantified using an HIV-1 p24 ELISA assay. The wild type SARS-CoV-2 (hCoV-19/Canada/ON-VIDO-01/2020, GISAID accession# EPI_ISL_425177) was propagated and produced in Vero cells (ATCC, CCL-81).

To investigate the infection ability of SCoV-2-SP-VPs, the same amount of each SCoV-2-SP-PV stock (as adjusted by p24 levels) were used to infect different target cells at 0.4 ⨯ 10^5^ cells per well (24 well plate) for 3 hrs and washed. After 48 or 72 hrs, the supernatants were collected and the viral infection rate was evaluated by measuring Gaussia luciferase (Gluc) activity. Briefly, 50ul of Coelenterazine substrate (Nanolight Technology) was added to 20ul of supernatant, mixed well and read in the luminometer (Promega, USA).

To evaluate the anti-SARS-CoV-2 SP-mediated entry activity of CHPV or suramin, various concentrations of herb extract or compound were directly added into target cells at different time points before or after infection, as indicated. After 3hrs of infection at 37°C, the cells were washed once to remove excessive residue viruses/compound and cultured in fresh medium. The anti-SARS-CoV-2 effects of CHPV or suramin were evaluated by measuring the Gluc activity or p24 levels in the supernatant infected cultures.

Efficacy of CHPV or suramin against SARS-CoV-2 (hCoV-19/Canada/ON-VIDO-01/2020, GISAID accession# EPI_ISL_425177) was evaluated in Vero cells. The Vero cells were seeded into 96-well plates and reached a confluency of 90% at the second day. Then each compound was diluted in assay medium (MEM with 1X penicillin-streptomycin) and added to the wells (100 ul/well), followed by adding 100 μL of SARS-CoV-2 at a MOI of 0.01, resulting in a final 1X drug concentrations. As positive controls, wells without drugs were infected with SARS-CoV-2 at the same MOI. Cells were maintained for 72 hrs and then, virus infection induced cytopathic effect (CPE) was monitored in each well.

### Binding Assay

The inhibitory effect of CHPV or suramin on the interaction of SP-ACE2 was tested with COVID-19 Spike-ACE2 binding assay kit. Briefly, 96-well plate was coated with recombinant SARS-CoV-2 Spike protein. CHPV or suramin was then added to the wells for 10 min followed by adding recombinant human ACE2 protein. After incubation for 3 hours, wells were washed three times and a goat anti-ACE2 antibody that binds to the Spike-ACE2 complex was added followed by applying the HRP-conjugated anti-goat IgG and 3,3’,5,5’-tetramethylbenzidine (TMB) substrate. The intensity of the yellow color is then measured at 450 nm.

### Western blot (WB) analyses

To detect cellular protein ACE2, SARS-CoV-2-SP, or SPΔC in transfected cells or SCoV-2-SP-VPs, transfected 293T_ACE2_ cells or VPs were lysed in RIPA buffer, and directly loaded into the 10 % SDS–PAGE gel and the presence of each protein was detected by WB with various corresponding antibodies.

## Results

### Generation of a SARS-CoV-2 SP-pseudotyped HIV-1-based entry system

We first established a sensitive SARS-CoV-2-SP-mediated virus entry system by co-transfecting SARS-CoV-2 SP, a HIV-based vector (ΔRI/ΔEnv/Gluc) in which viral reverse transcriptase/integrase deleted/envelope gene partially deleted and encoded a Gaussia luciferase gene in the *nef* position [34], and a packaging plasmid (pCMVΔR8.2) in HEK293T cells (Fig. 1B). The Gaussia luciferase (Gluc) is a bioluminescent enzyme that can be secreted into the media, enabling the analysis of viral expression by direct measurement of Gluc activity in the supernatant. To do this, we have constructed a full length SP (SARS-CoV-2-SP) and the C-terminal 17 amino acid (aa) deletion SP (SARS-CoV-2-SPΔC) expressing plasmids since previous studies have reported that a carboxyl-terminal truncation of 17 amino acids of SARS SP substantially increased SARS SP-mediated cell-to-cell fusion [37].

**Figure 1.**
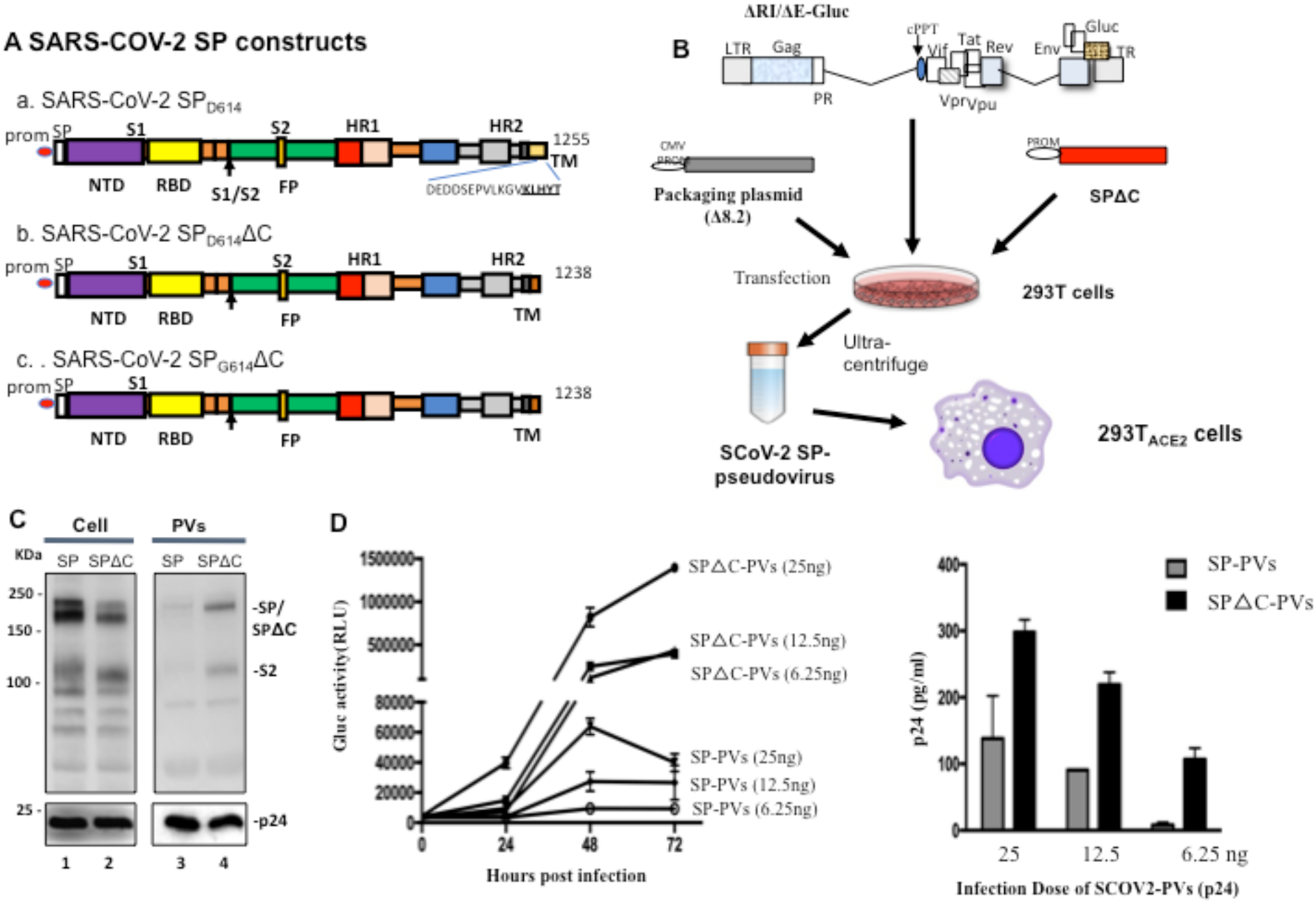
Generation of a SARS-COV2-SP-pseudotyped lentiviruse particles (SCoV-2-SP-PVs). A) Schematic representation of SARS-CoV-2SP, SARS-CoV-2SPΔC, and SARS-CoV-2SP_G614_ΔC expressing plasmids. B) Schematic representation of plasmids and and procedures for production of SARS-COV2-SP-pseudotyped lentivirus particles (SCoV-2-SP-PVs). C) Detection of SARS-CoV-2 SPs and HIV p24 protein expression in transfected 293T cells and viral particles by Western blot (WB) with anti-SP or anti-p24 antibodies. D) Different amounts of SCoV-2-SP-PVs and SCoV-2-SPΔC-PVs virions (adjusted by p24) were used to infect 293T_ACE2_ cells. At different time intervels, the Gaussia Luciferase activity (Gluc) (left panel) and PVs-associated p24 (at 72 hrs) in supernatants was measured.

To examine the expression and incorporation of SARS-CoV-2 SPs and SPΔC in the cells and the SARS-COV-2-SP pseudotyped viruses (SP-PVs) and SARS-COV-2-SPΔC pseudotyped viruses (SPΔC-PVs), lysates of both virus-producing cells and pseudotyped viruses were analyzed by SDS-PAGE and WB with a mouse anti-SP antibody, as indicated in Fig. 1C. As expected, the HIV capsid Gagp24 protein was detected in all of the cell lysates and the pelleted SP-PVs and SPΔC-PVs pseudoviruses (PVs) by rabbit anti-p24 antibodies (Fig. 1C, lower panel). The SARS-CoV-2 SP including S1/S2 were clearly detected in both SARS-CoV-2-SPs and SARS-CoV-2-SPΔC-transfected cells (Fig. 1C, lane 1). Interestingly our results revealed that virus-incorporation level of SARS-CoV-2-SPΔC were significantly higher than that of SARS-CoV-2-SP (Fig. 1C, compare lane 4 to 3), To test the infectivity of generated pseudoviruses, we infected 293T-ACE2 cells with serial diluted amounts of pseudoviruses (25, 12.5, 6.25ng of p24) of SP-PVs or SPΔC-PVs for 3 hrs. The Gluc activities or Gagp24 of supernatants from infected cells were measured at 24h, 48h or 72h post infection. The results showed that both SP-PVs and SPΔC-PVs can infect 293T-ACE2 cells and induce an increase of Gluc activity in the supernatants in a dose dependent manner (Fig.1D, left panel). As expected, the infectivity of SPΔC-PVs was significantly higher than that of SP-PVs. The infection of pseudoviruses in 293T_ACE2_ cells was further confirmed by detection of the HIVp24 levels in the supernatants of infected cells through ELISA assay (Fig.1D, right panel).

To test whether the infection is ACE2-dependent, we infected various cell lines, including HEK293T, 293T_ACE2_ and VeroE6 with SP-PVs and SPΔC-PVs, respectively. The results showed that these pseudoviruses were only able to efficiently infect 293T_ACE2_ cells, and not HEK293T or VeroE6 cells (Fig. 2A). In parallel, we only detected high level expression of the SARS-CoV-2 receptor ACE2 in 293T_ACE2_ cells, but not in the 293T or Vero E6 cells (Fig. 2B).

**Figure 2.**
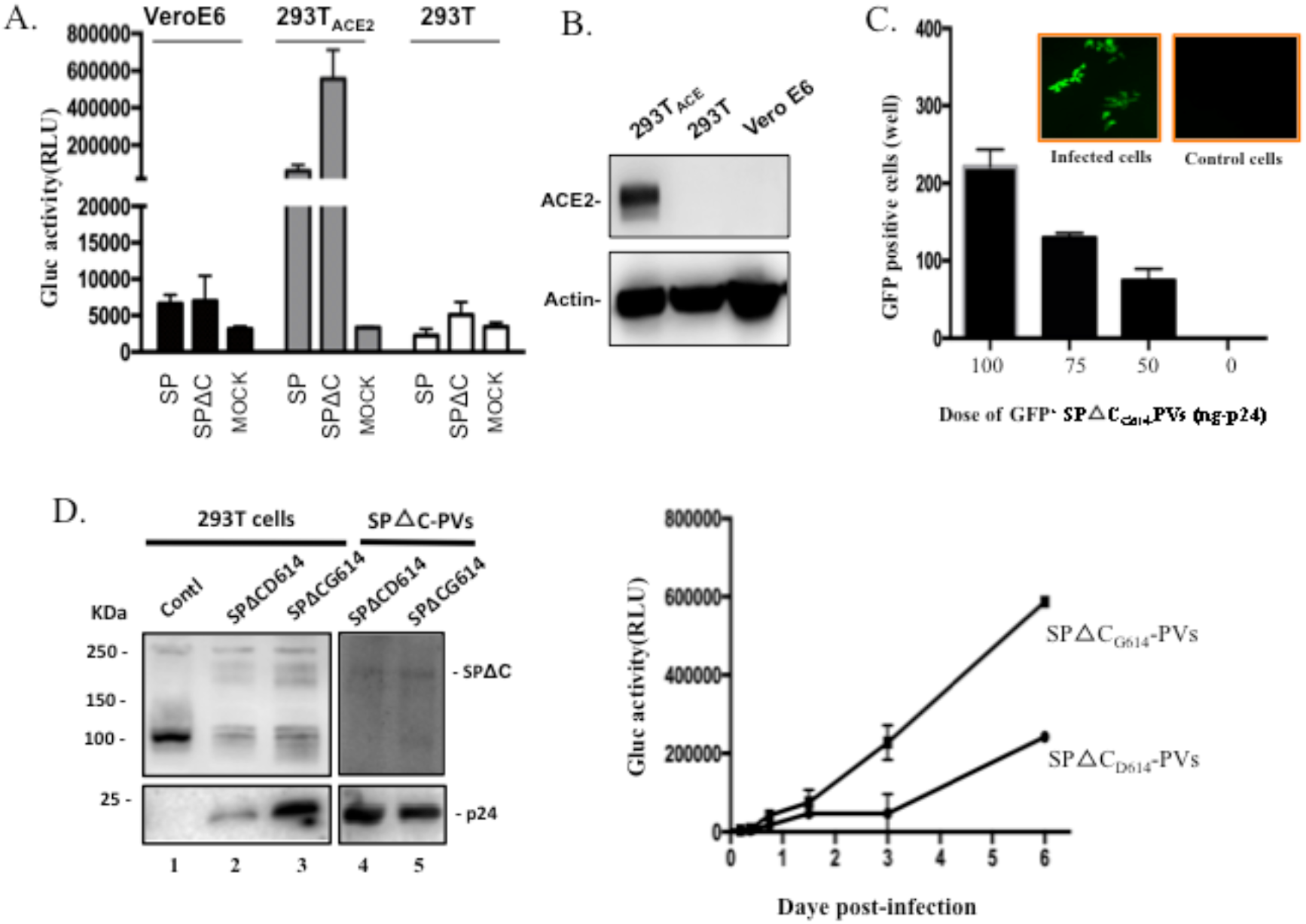
SARS-CoV-2 SP-PVs’s infection in different cell lines and SARS-CoV-2 SP_G614_ variant exhibited stronger virus entry. A) 293T, 293T_ACE2_ and Vero-E6 cells were infected by equal amounts of SARS-CoV-2SP-, SARS-CoV-2SPΔC-pseudotyped viruses. At 48 hrs pi, the Gluc activity in supernatants was measured. B) the expression of SARS-CoV-2SP receptor, ACE2, in 293T, 293T_ACE2_ and Vero-E6 cells detected by WB with anti-ACE2 antibodies. C) The SPΔC_G614_-GFP^+^PVs were produced with 293T cells and used to infect 293T_ACE2_ cells in 96-well plate After 48 hrs pi, GFP-positive cells (per well) were counted and photographed by fluorescence microscope (on the top of the panel). D) Detection of SARS-CoV-2 SPΔC, SPΔC_G614_ and HIV p24 protein expression in transfected 293T cells and viral particles by WB. E) Infectivity comparison of SPΔC-PVs and SPΔC_G614_-PVs in 293T_ACE2_ cells. Equal amounts of SPΔC_D614_-PVs and SPΔC_G614_-PVs virions (adjusted by p24 level) were used to infect 293T_ACE2_ cells. At different days post-infection (pi), Gluc activity in supernatants was measured.

**Figure 3.**
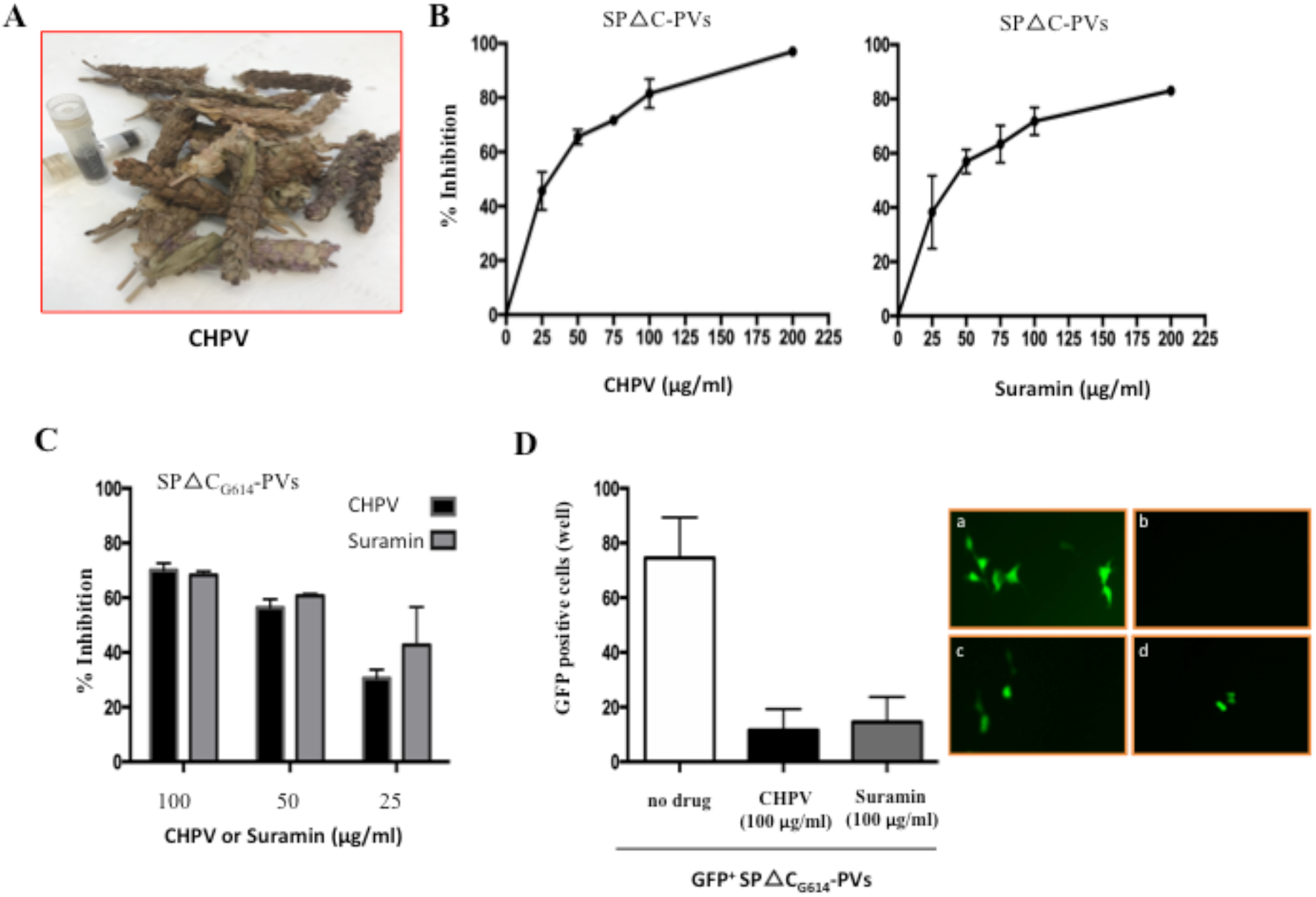
SARS-CoV-2-SP-PV’s infection was efficiently blocked by CHPV and suramin. A) Images of the dried Prunella Vulgaris flowers and its water extract (CHPV). B) Dose -response anti-SARS-CoV-2 analysis by Gluc activity for CHPV or suramin. 293T_ACE2_ cells were infected by equal amounts of SARS-CoV-2SPΔC-pseudotyped viruses in the presence of different dose of CHPV or suramin. At 48 hrs pi, the Gluc activity in supernatants was measured. (% inhibition = 100 ⨯ [1 - (Gluc value in presence of drug)/(Gluc value in absence of drug)). C) Infection inhibition of CHPV or suramin on SARS-CoV-2-SPΔC_G614_-PVs in 293T_ACE2_ cells. Equal amounts of SCoV-2-SPΔC_G614_-PVs (adjusted by p24 level) were used to infect 293T_ACE2_ cells in presence of different concentrations of CHPV or suramin, in indicated at bottom of the panel. At 48 hrs pi, Gluc activity in supernatants was measured and present as % inhibition. Means ± SD were calculated from duplicate experiments. D) 293T_ACE2_ cells in 96-well plate were infected with SPΔC_G614_-GFP^+^ PVs. After 48 hrs pi, GFP-positive cells (per well) were counted (left panel) and photographed by fluorescence microscope (right panel, a. Without drugs; b. Without infection; c. In the presence of CHPV (100 μg/ml); d. In the presence of suramin (100 μg/ml).

In addition, we have generated a GFP^+^ SARS-CoV-2-SP-mediated virus entry system by cotransfecting SARS-CoV-2 SPΔC_G614_, a lentiviral vector that expressing GFP, and the pCMVΔR8.2 in HEK293T cells and produced SPΔC_G614_-PVs expressing GFP (GFP^+^ SPΔC_G614_-PVs). After 293T_ACE2_ cells were infected with GFP^+^ SPΔC_G614_-PVs, the GFP positive 293T_ACE2_ cells were clearly detected under fluorescent microscopy (Fig. 2C)

### 2. SARS-CoV-2 SP G614 variant exhibited stronger receptor association and virus entry

Recent sequence analyses revealed a SP mutation, Aspartic acid (D) changed to Glycine (G) at aa position 614, was found in high frequency (65% to 70%) in April to May of 2020, indicating a transmission advantage to D614 [15]. In this study, we have also generated constructs to express SCoV-2-SPΔC_G614_ (Fig. 1A,c) and compared its virus entry ability with SCoV-2-SPΔC (SPΔC_D614_). Our results showed that SCoV-2-SPΔC_G614_ was incorporated into pseudotyped viruses similar to SCoV-2-SPΔC_D614_ (Fig. 2D, compare lane 5 to lane 4). However, the SARS-CoV2-SPΔC_G614_-pseudotyped viral particles (SPΔC_G614_-PVs) mediated approximately 3-fold higher infection than that of SPΔC_D614_-PVs (Fig. 2E), suggesting that the SP_G614_ mutation increases SP-mediated viral entry.

### Evaluation of CHPV and Suramin for blocking SARS-CoV2-SP-mediated virus entry

Next we tested whether CHPV (Fig. 3A) and suramin could block SARS-CoV2 SP-mediated virus entry of 293TACE2 cells. Briefly, 293TACE2 cells were infected with SPΔC-PVs in the presence of different concentrations (25, 50,75, 100 and 200ug/ml) of CHPV (Fig. 3B) or suramin (Fig. 3C), respectively. After 3 hour of infection, the infected cells were washed to remove the viruses and compounds and cultured with fresh medium. At 48 hrs post-infection, the supernatants were collected and the virus-produced Gluc activities were measured for monitoring the infection levels. Consistent with our previous observation [26], we did not detect any CHPV-induced toxic effect on the cells for 3hs exposure, nor for Suramin. Significantly, both CHPV and suramin were able to inhibit SARS-CoV-2-SP-pseudotyped virus infection. The half maximal inhibitory concentration (IC50) of CHPV was 30 ug/ml (Fig.3. B, left panel, while IC50 of Suramin was about 40 ug/ml (Fig.3. B, right panel). The inhibitory effect of CHPV and suramin on a SP mutant pseudotyped virus (SPΔC_G614_-PVs) infection was also tested, and results show that SPΔC_G614_-PVs infection is also susceptible to CHPV and suramin (Fig. 3C). Furthermore, the SARS-CoV-2-SPΔC_G614_ psedotyped GFP+ virus infection was tested in the presence of the two compounds and results showed that the psedotyped GFP+ virus infection was efficiently inhibited by the presence of CHPV and Suramin (Fig. 3D). All of these results demonstrate that both CHPV and suramin exhibit strong inhibitory effect on both SPΔC_D614_-PVs and SPΔC_G614_-PVs infection.

### Mechanistic analyses of actions of CHPV and Suramin against SARS-CoV2-SP-mediated virus entry

To gain more insight into the mechanism of how CHPV and Suramin are targeting SARS-CoV-2 SP-PVs infection, each of the drugs (100 μg/ml) was added to 293T_ACE2_ cells at various time points during the infection, as indicated in Figure 4. After 48 hrs of infection, the supernatants were collected and measured for virus-expressed GLuc activity. Results showed that a strong inhibitory effect was achieved when cells were pretreated with CHPV or Suramin one hour before infection or when the compounds were present simultaneously with SP-PVs (Fig. 4A and B). Interestingly, even when drug was added at one hr post-infection, CHPV still exhibited nearly 70% inhibition on SPΔC_G614_-PVs infection (Fig. 4A), while for Suramin, a lower inhibitory effect (about 30% inhibition) was also observed (Fig. 4B). When CHPV or Suramin was added to culture after 3 hrs of infection, no inhibitory activity on viral infection was observed (Fig. 4A and B). These results suggest that both CHPV and suramin act on the entry step of SPΔC_G614_*-*PVs infection.

**Figure 4.**
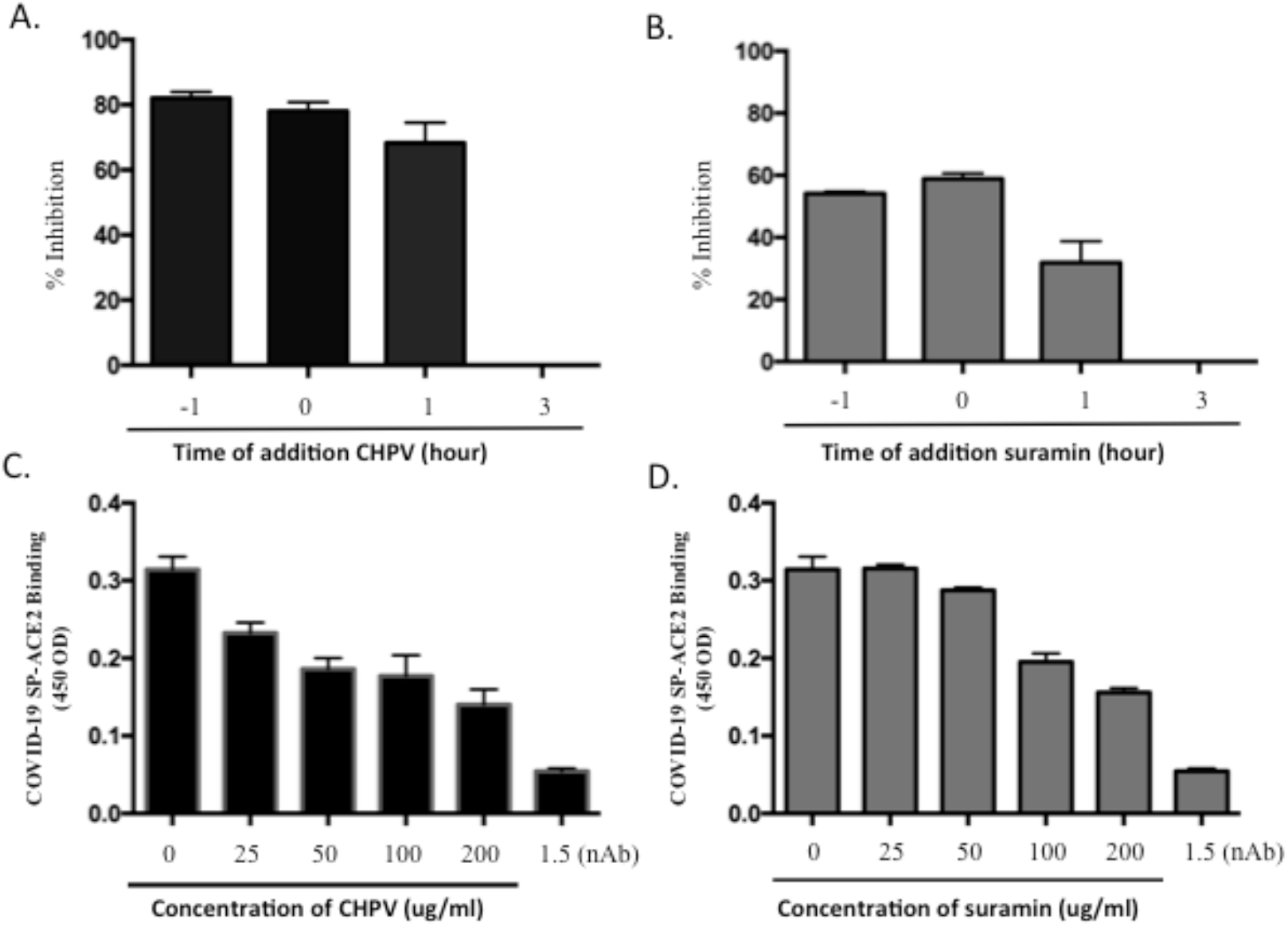
Characterization of the mechanisms of CHPV and suramin for their anti-SARS-COV-2-SP action. A) Time-dependent inhibition of SPΔC_G614_-PVs infection mediated by CHPV or suramin. CHPV (100 μg/mL) or suramin (100 μg/mL) was added at 1 hr prior to infection, during infection (0 hr), and at 1 hr, and 3 hr pi. The positive controls (PC) were 293T_ACE2_ cells infected with SPΔC_G614_-PVs in the absence of compounds. At 3 hrs pi, all of the cell cultures were replaced with fresh DMEM and cultured for 48 hrs. Then, the Gluc activity was monitored in the supernatant, and the data are shown as a percentage of inhibition (%). B) inhibitory effect of CHPV or suramin on SARS-CoV2-SP/ACE2 binding by ELISA as described in materials and methods. nAB: anti-COVID-19 neutralizing antibody (SAD-S35). The results are the mean ± SD of duplicate samples, and the data are representative of results obtained in two independent experiments.

To further determine whether CHPV or suramin is targeting the interaction of SARS-CoV2-SP and its receptor, ACE2,, we used an *in vitro* SARS-CoV2-SP/ACE2 binding ELISA assay, as described in Materials and Methods. Additionally, an anti-COVID-19 neutralizing antibody (SAD-S35) [38] was included in parallel. Results revealed that the presence of either CHPV or suramin was able to specifically target and significantly reduce the SARS-CoV2-SP-ACE2 interaction (Fig. 4C and D). It should be noted that the neutralizing antibody (SAD-S35) also showed a strong inhibition on SARS-CoV2-SP/ACE2 interaction (Fig. 4C and D).

### Combination of CHPV and anti-SARS-CoV-2 neutralizing antibody (SAD-S35) mediated more potent blocking effect against SARS-CoV2-SP-PVs

As described above, both CHPV and Suramin can inhibit SARS-CoV2-SP/ACE2 interaction and SP-PVs infection. We also tested whether the combination of two compounds could mediate a stronger anti-SARS-CoV-2 activity. Thus, we infected 293T_ACE2_ cells with SPΔC_G614_-PVs in the presence of a cocktail of CHPV/Suramin (25 μg/mL per compound), or CHPV (50 μg/mL) or Suramin (50 μg/mL) alone. The results showed that in the presence of a cocktail of CHPV/Suramin, SPΔC_G614_-PVs was inhibited to 78%, while in the presence of CHPV or suramin alone, inhibition rate was 65% or 40% (Fig. 5A). These results suggest that a combination of these two compounds may be able to achieve more efficient inhibition against SARS-CoV-2 infection.

**Figure 5.**
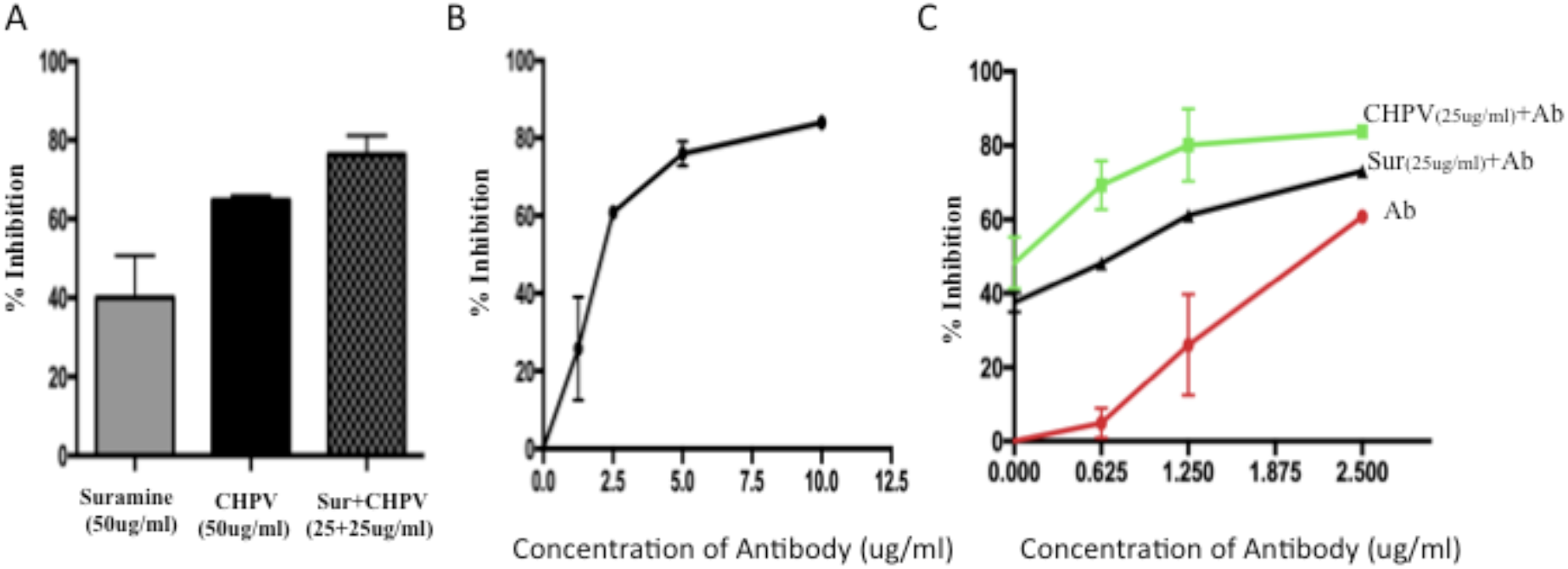
Enhanced inhibitory effects mediated by combination of CHPV and suramin with neutralizing antibody (SAD-S35). A) 293T_ACE2_ cells were infected with SPΔC_G614_-PVs in presence of CHPV (50μg/ml) or suramin (50μg/ml) alone or a mix of CHPV and suramin (each with 25μg/ml). After 3 hrs of infection, cells were washed and add fresh medium for 48 hrs. Then the supernatants were collected and Gluc activity in the supernatant was measured and present as % inhibition. B) Inhibitory effect of nAb SAD-S35 on SPΔC_G614_-PVs infection. 293T_ACE2_ cells were infected with SPΔC_G614_-PVs in the presence of serially diluted SAD-S35 (1.25 to 10 μg/ml) for 3 hrs. Then infected cells were cultured in fresh medium. At 48 hrs pi., the supernatants were collected and measured for Gluc activities and presented as % inhibition. C) 293T_ACE2_ cells were infected with SPΔC_G614_-PVs in the presence of serially diluted SAD-S35 (0.625 to 2.5 μg/ml) alone or mixed with CHPV (25 μg/mL) or Suramin (25 μg/mL) for 3 hrs and the infected cells were cultured in fresh medium. At 48 hrs pi., the Gluc activities in the supernatants were measured and presented as % inhibition. The results are the mean ± SD of duplicate samples, and the data are representative of results obtained in two independent experiments.

The anti-SARS-CoV-2 neutralizing antibody (SAD-S35) was also tested and showed a does-dependent neutralizing activity against SPΔC_G614_-PVs with an IC50 of 2.4 μg/mL (Fig. 5B). Next, we sought to determine whether the combination of CHPV or Suramin with SAD-S35 could mediate a stronger anti-SARS-CoV-2 activity. Thus, serially diluted SAD-S35 (0.625 to 2.5 μg/ml) was mixed with CHPV (25 μg/mL) or Suramin (25 μg/mL) and added to the 293T_ACE2_ cells with SPΔC_G614_-PVs simultaneously. In parallel, same concentrations of SAD-S35 alone were used for comparison. The results show that 1.25 μg/ml of SAD-S35 alone only resulted in an approximately 25% decrease of infection. However, nearly 80% inhibitory effect was achieved when the same concentration of SAD-S35 was combined with CHPV (25μg/ml), or approximately 60% inhibitory effect was achieved when combined with suramin (25μg/ml), while CHPV or suramin alone only mediated 50% or 38% inhibition, respectively (Fig. 5C). All together, the results clearly indicate that a combination of CHPV or suramin with SAD-S35 is able to more potently block SARS-CoV2 infection. By including a low dose of nAb, the amounts of CHPV or Suramin needed to achieve highly effective inhibition of SARS-CoV2 infection can be reduced.

### Inhibitory effect of CHPV and Suramin on SARS-CoV-2 virus infection

Given that both CHPV and suramin are able to block the SARS-CoV2-SP pseudovirus entry, we next tested whether these two drugs could block wild type SARS-CoV2 virus infection and virus-induced cytopathic effect in Vero cells. The wild type SARS-CoV-2 virus (hCoV-19/Canada/ON-VIDO-01/2020) was used to infect Vero cells in the presence of different concentrations of CHPV or Suramin. Briefly, Vero cells were infected with SARS-CoV-2 (MOI of 0.01) in the presence of different concentrations of CHPV or Suramin. After 72 hrs post-infection, as indicated (Fig. 6), the SARS-CoV-2-induced cytopathic effects in Vero cells were monitored. Results showed that SARS-CoV-2 infection causes dramatic cytopathic effect (CPE) in Vero cells after 72 hrs post-infection, with cells displaying 100% CPE. Remarkably, in the presence of CHPV or suramin (at 50 to 125 μg/ml), the SARS-CoV-2-induced cytopathic effect (CPE) was significantly or completely inhibited in Vero cells. These results provide strong evidence that the presence of CHPV or Suramin is able to inhibit SARS-COV-2 infection.

**Figure 6.**
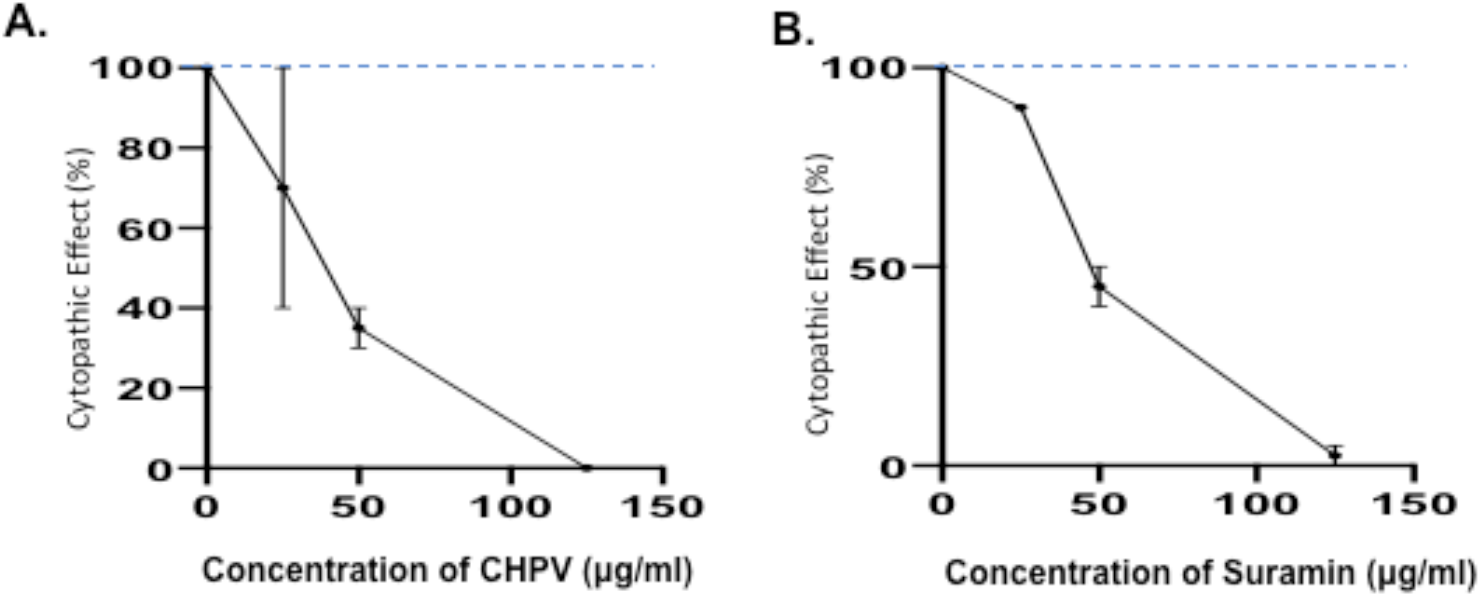
Inhibitory effect of CHPV and Suramin on SARS-CoV-2 infection-induced cytopathic effects. Vero cells were infected with a wild type *SARS-CoV-2* virus (hCoV-19/Canada/ON-VIDO-01/2020) in the presence or absence of different concentrations of CHPV and Suramin. After 72 hrs pi., the SARS-CoV-2 infection-induced cytopathic effects in Vero cells were monitored. Error bars represent variation between triplicate samples, and the data of (A) and (B) are representative of results obtained in two independent experiments.

## Discussion

Because SARS-COV-2 is classified as an aerosol biosafety level 3 (BSL-3) pathogen, the study of SARS-COV-2 infection and the investigation of different anti-SARS-COV-2 compounds required highly restricted BSL-3 containment. This condition has significantly limited the SARS-COV-2-related research activities in microbiology laboratories. In this study, we established a highly sensitive SARS-COV-2-SP psedotyped HIV-based entry system, which encodes a Gaussia luciferase (Gluc) gene as a reporter (Fig. 1B). Since Gluc can be secreted into the supernatant after being expressed in the infected cells, it is very sensitive and convenient for evaluating the level of SARS-CoV2-SP-mediated virus entry and may be used for anti-SARS-CoV2-SP compound screening in a BSL-2 environment.

Previous studies have revealed that the cytoplasmic tail (CT) of SARS SP contains a dibasic motif (KxHxx) that constitutes for an endoplasmic reticulum (ER) retrieval signal which retains the full-length SARS-S protein in the lumen of the ER-Golgi intermediate compartment (ERGIC) [39,40]. Deletion of 17 aa at the carboxyl-terminal in the CT of SARS SP was able to increase SP transported to the surface of cells and substantially increased SARS SP-mediated cell-to-cell fusion [37]. In the SARS-CoV2-SP, there is also a dibasic motif (KxHxx) present in the CT (Fg 1A). In order to increase SARS-CoV2-SP incorporation into pseudovirions, we have deleted 17 aa at the CT of SARS-CoV2 SP and generated a SARS-CoV2 SPΔC expressor plasmid (Fig. 1A, b). Indeed, our data showed that a significantly higher level of SCoV2 SPΔC protein was present in the pseudovirus (Fig. 1C), and induced remarkably efficient infection in 293T_ACE2_ cells (Fig. 1D). This observation clearly indicate that the dibasic motif in SARS-COV-2 SP is functional and a deletion of 17 amino acids substantially increased incorporation of SP into SARS-CoV2-SP-PVs and enhance its infectivity. This information is also important for improving the design of SARS-CoV2-SP-based vaccine strategies.

Recent sequencing analyses found a SARS-CoV2 SP mutation, Aspartic acid (D) changed to Glycine (G) at aa position 614 in the S1 domain which was dominantly detected in April to May of 2020 isolates, indicating a transmission advantage over original SP D614 [15]. The following studies showed that SARS-CoV2_G614_ SP mutant MLV pseudotyped viruses infected ACE2-expressing cells markedly more efficiently than those with SARS-CoV2_D614_ [16,41]. Consistently, we also observed the SARS-CoV2_G614_ΔC-pseudotyped lentiviral particles enhanced the pseudotyped virus entry compared to the SP_D614_ΔC-PVs (Fig. 2E).

By using this SCoV2-SP-pseudovirus system, we have provided evidence for the first time that the CHPV can efficiently prevent infections mediated by both SARS-CoV2-SP_D614_ and -SP_G614_ pseudovirus infection in 293T_ACE2_ cells and significantly block the infection of wildtype SARS-COV-2 in Vero cells. We also revealed that CHPV blocks the entry of virus by directly interrupting the interaction of SARS-CoV2-SP and ACE2 receptor by *in vitro* ELISA assay (Fig. 4). Interestingly, the presence of CHPV at one hour post-infection is still able to efficiently inhibit SARS-Cov2-SP pseudovirus infection (Fig. 4), suggesting that CHPV may not only target SP/ACE2 binding, but may also act on the following fusion step(s). Overall, our results provide convincing evidence for CHPV as a potential blocking agent against SARS-COV-2 infection. In agreement with a recent study [33] that suramin can inhibit SARS-COV-2 virus infection, we further provide evidence that suramin is able to directly block SARS-CoV-2 SP-ACE2 interaction (Fig. 4D) and different SARS-CoV-2 SP variants mediated virus entry (Fig 3, 4, and 5).

Another interesting observation in this study is that the combination of CHPV or suramin with anti-SARS-COV-2 neutralizing antibody (nAb) could enhance their anti-SARS-COV-2 activity. The nAb has great potential to be used as a preventing agent in blocking SARS-COV-2 infection [42]. However, one disadvantage of using nAb as an anti-SARS-COV-2 agent is its source limitation. Therefore, the finding of the synergistic effect of a combination of nAb with other agents, such as CHPV or Suramin is beneficial for (i) similar efficiencies would be achieved by using reduced amounts of antibody and CHPV or Suramin, (ii) the combination of nAb and CHPV/suramin will reduce the likelihood of viral resistance. Whether these enhanced effects might be due to a combined effect through their different binding mechanisms still needs to be investigated.

The effectiveness of CHPV and/or suramin against SARS-COV-2 infection *in vivo* remains to be investigated. Our findings could be further validated in an appropriate animal model and clinical trials for prevention of COVID-19. Since SARS-COV-2 infection initiates in the respiratory tract [43], the use of CHPV and/or Suramin as nasopharynx agents (Nasal spray) to prevent initial SARS-COV-2 infection and transmission in the respiratory tract will be a particularly attractive strategy, and will require further efficacy studies. Overall, we demonstrated that CHPV and suramin possess an anti-SARS-COV-2 entry inhibitor activity and functions at least partially by interrupting SARS-COV-2 SP binding to its receptor. Additional *in vivo* safety and protection studies will facilitate its application as an option to help control the ongoing SARS-CoV-2 pandemic.

## Acknowledgements

Dr. X-j Yao holds a Manitoba Research Chair Award from the Research Manitoba. O.T.A is the recipient of PhD scholarship from the Research Manitoba/Manitoba Institute of Child Health. This work was supported by Canadian 2019 Novel Coronavirus (COVID-19) Rapid Research Funding (OV5-170710) by Canadian Institute of Health Research (CIHR) and Research Manitoba.

